# Social information use for spatial decision in *Zootoca vivipara*

**DOI:** 10.1101/2021.10.08.463627

**Authors:** M. Brevet, S. Jacob, A. Rutschmann, M. Richard, J. Cote, J. Clobert

## Abstract

Movements of individuals are conditioned by information acquisition coming from either personal or social sources. Yet, little is known about the processes used by individuals to make movement decisions when facing multiple sources of social information simultaneously. This study aimed to test experimentally how social information from multiple sources is used to make movement decisions, and whether a contrast in this information allows individuals to orientate in space. We used common lizards (*Zootoca vivipara*) in a replicated experimental setting: one focal individual received information from two other individuals coming from peripheral environments, before being given the opportunity to relocate in one or another of the peripheral environments.

Our analyses revealed that the behavior of informants, their mother’s morphology, as well as the quality of informants’ environment, affected movement decisions: the probability to relocate from the focal area increased when informants displayed traits associated with low resources (no food intake, poor maternal condition) or high competition (high activity). The physical condition of individuals also mediated the use of social information about food intake, with a match between resource availability in informants and personal condition. Conversely, spatial orientation was not affected by the contrast of phenotype between informants nor by spatial variability in resource availability.

This study highlights that multiple social information sources can be used for movement decisions, likely because these information sources reflect the quality of the surrounding environment (e.g., competition level or resources availability). It also emphasizes that social information use for movement is conditioned by individual phenotype.

## Introduction

Information acquisition is central for an individual to assess the quality of its environment and to take appropriate decisions to feed, survive and reproduce (Dall et al. 2005, Schmidt et al. 2010). Information can either be obtained via personal interactions with the environment (i.e., personal information, Dall et al. 2005) or via social interactions (i.e., social information, Dall et al. 2005, Schmidt et al. 2010). Social information is acquired through the observation of conspecific’s or heterospecific’s detectable traits (e.g., behavior, performance, body condition, odors; Moreira et al. 2008, Clobert et al. 2009) and can inform individuals about both abiotic and biotic characteristics of the local and distant environments such as breeding habitat quality (Doligez et al. 2002 and 2004) or conspecific density and resource availability (Endriss et al. 2018). Social information can be intentionally transmitted by informant individuals through signals such as calls or territorial marking (Johnson 1973, Macedonia et al. 1993) but may also be inadvertently conveyed by cues (Schmidt et al, 2010), as it is the case for breeding habitat quality, informed by reproduction performances in some bird species (Doligez et al. 2002, 2004).

Social information has long been recognized to be key in organisms’ decisions to move through their environment and is notably known to influence the optimization of spatial decision making for microhabitat use (e.g., Moreira et al. 2008), habitat selection (Doligez et al. 2002, 2004) or dispersal (Cote and Clobert 2007, Jacob et al. 2015). In spatially heterogeneous environments, social information is expected to be especially useful for any movement decision. Indeed, in such environments, the social information is made of a mosaic of cues or signals carried by either local inhabitants or immigrants, respectively informing on the specificities of close and distant habitats (Cote and Clobert 2007, Jacob et al. 2015). With increasing heterogeneity, environmental predictability is expected to decrease and socially acquired information can increase an individual knowledge of the general environment, hence reducing the probability of erroneous decisions due to environmental uncertainty (Dall et al. 2005, Riotte-Lambert et al. 2020). In other words, informed individuals are more likely to make the right movement decisions (i.e., the one that maximizes their fitness) if they acquire knowledge about the suitability of surrounding habitats throughout social information, compared to when relying on their local prospects only, especially if their movement abilities are restricted.

Yet, little is known about the use of social information for movements decision when multiple sources of social information are simultaneously accessible to an individual (i.e., multiple conspecifics reflecting different surrounding habitats). More specifically, two important questions have to be addressed. First, when multiple sources of information are available, how information sources are used to decide whether the individual should relocate or not? One may expect that the averaged information on surrounding habitats should prevail (Hyp. 1.a: Social information synthesis, figure 1): an individual would optimize its movement by synthesizing all sources of information available (i.e., each social cue or signal among sources), whatever the quality and origin of the cue or signal, to get a global idea of the amount of resource and level of intraspecific competition in the vicinity (Stamps 2001, Clobert et al. 2004, Bowler and Benton 2005). The use of information on surrounding habitats could also depend on the phenotype of the informed individual, which would adjust movement decision to its condition and relocate only when necessary (Hyp. 1.b: Phenotypic-dependent social information synthesis, figure 1). Many examples in the literature illustrate such phenotype-dependent use of social information, with dependence on personality (Smit and van Oers 2019, Morinay et al. 2020), age and success (Parejo et al. 2007), or body condition (Cote and Clobert 2007). One may also expect individuals to use the contrast between available information sources (Hyp 1.c: Contrasted social information use, figure 1): by comparing the concordance or discordance of social cues or signals between sources, individuals might be able to assess information reliability or environmental variability in the vicinity. The importance of conflictual information for movement decisions has already been observed for conflicts between personal and social information (e.g., Cronin 2013, Winandy et al. 2020), with prioritization of personal information in case of conflicts (Kendal 2009).

**Figure 1:**
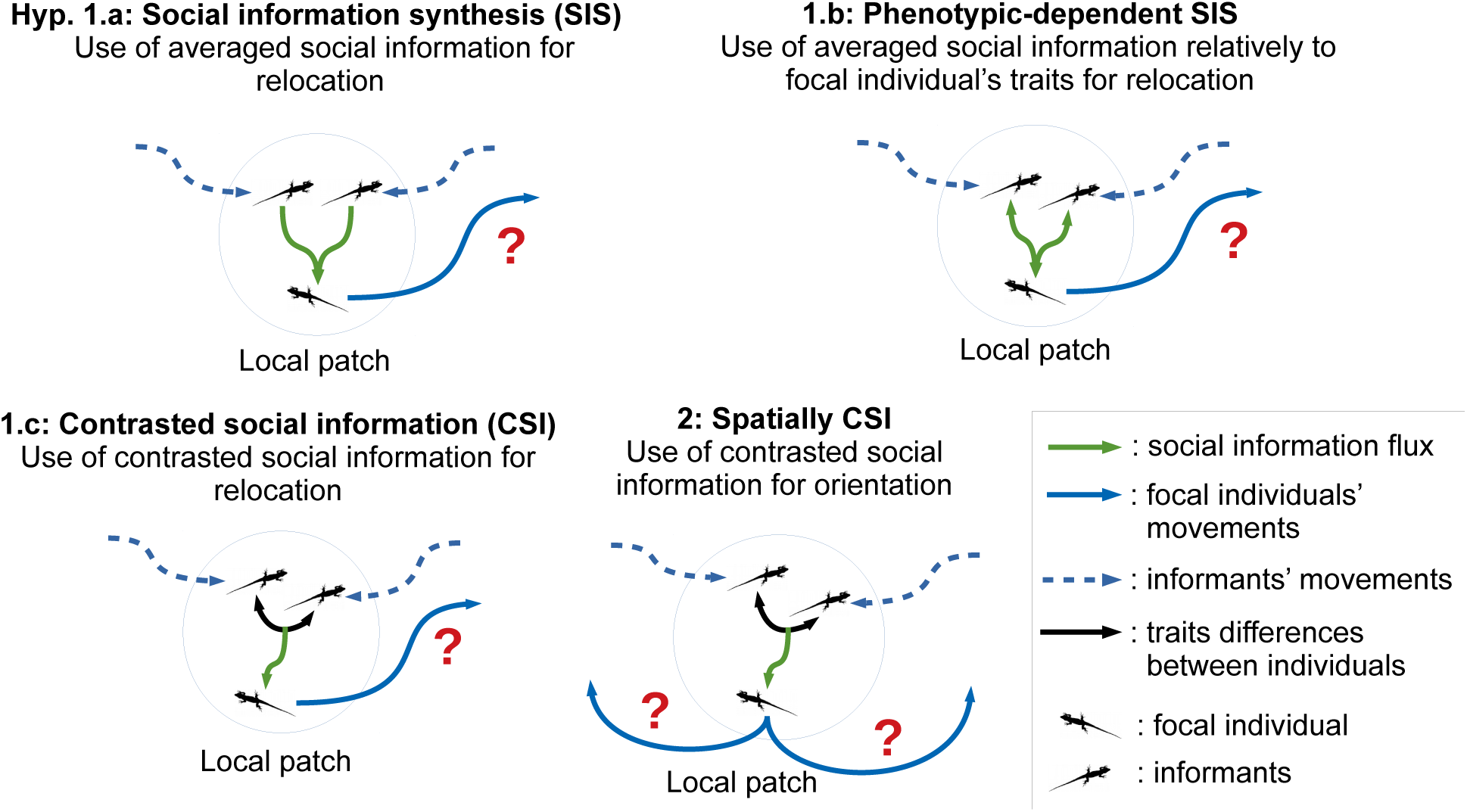
Experimental aims. These schemes present the different hypotheses that we investigated through our experimental design. We hypothesized that relocation would depend on a synthesis of all social information on surrounding habitats. Such use of multiple sources of social information might consist in using an averaged value of multiple information sources (Hyp. 1.a), potentially mediated by focal individual’s phenotype (Hyp. 1.b). Conversely, the use of social information could consist in using the contrast between information sources as an indicator of information variability/reliability (Hyp. 1.c). Finally, we hypothesized that the existing contrast between information sources could allow the focal individual to orientate in space in order to join the best of the surrounding habitats (Hyp. 2).

The second question lies in the spatial integration of information gathered from multiple sources coming from different locations: how does an individual use multiple sources of information to orientate, and therefore to choose a specific destination of relocation between alternative habitats? One likely hypothesis (Hyp. 2: Spatially contrasted social information use, figure 1) is that differences in information between multiple sources originating from different habitats allow spatial orientation for the information receivers. Such differences in information could indeed inform the individual on the direction of habitats with a higher fitness expectancy, since the surrounding habitats are possibly associated with different social information quality as a function of their fitness expectancy (Schmidt et al. 2010).

To investigate these questions, we used the common lizard (*Zootoca vivipara*, Jacquin 1787) as a model species. This lizard is known to use social information in different contexts, and notably to acquire information about the dispersal status of conspecifics (Aragon 2006 b.), the reproductive strategy and aggressiveness of other females from their ventral coloration (Vercken et al. 2012), or the population density in the surrounding habitats through immigrants (Cote and Clobert 2007, Cote et al. 2008 a.). In this species, the use of personal and social information is also known to occur immediately after birth and to shape natal dispersal decisions (Clobert et al. 2012, Cote and Clobert 2012).

Here, we tested how common lizards use social information from two contrasted habitats, varying in food availability (present or not), to take decisions of relocation from their local area. We also tested whether such information could influence movements’ orientation when relocation occurred. To do so, we placed juvenile lizards (neonates from 2 to 4 days, born in our facilities from caught gravid females) in a three-chamber system (figure S1), where an information receiver (referred to as the focal individual further on) was confronted with two individuals coming from independent chambers and carrying contrasting information (referred to as informants further on). One informant came from an experimental environment where food was provided, while the other came from an experimental environment where food was absent. We generated all possible combinations of informants’ sexes introduced to the focal individual and measured multiple phenotypic traits that might convey information about informants and their habitats. We hypothesized that informants could transmit social information through their phenotype (e.g., behaviors, feeding, age, sex, or morphology) but also through their mothers’ condition (i.e., maternal effects, Bernardo 1996). These phenotypic traits are known to be related with short term sexual and resource competition contexts (Massot 1992, Léna et al. 1998, Galliard et al. 2005 a. for sex and morphology; Lecomte 1994, Cote et al. 2008 a. for behaviors), or with the long-term environmental context through maternal condition effects (Sorci and Clobert 1997, Uller and Olsson 2005, Mugabo et al. 2011). Note here that these phenotypic traits do not only inform about the competition in the standardized experimental setup, but possibly also about the competition level present in the habitat of origin of the individuals.

We expected relocation of the focal individual to vary with its phenotypic traits (Phenotype-dependence of movement) as already shown for body condition (Cote and Clobert 2007), sex (Galliard et al. 2005 a., Aragon et al. 2006 b.), age (Massot 1992, Léna et al. 1998), behaviors (Cote et al. 2010) or maternal body condition (kin competition, Léna et al. 1998, Meylan et al. 2002). Using this set-up, we tested whether the focal individuals could use social information from conspecifics to decide to relocate (or not) from their initial area. More specifically, we expected focal individuals to increase their movements when information about high quality surrounding habitats is on average provided (Hyp. 1.a, figure 1): low competition for resources (i.e., poorly active informants with poor physical conditions); long-term quality of the habitat (i.e., informants’ mothers with good physical conditions); low sexual competition (i.e., low male informants’ number) or sufficient resources availability (i.e., well-fed informants). The use of social information to adjust movement decisions might furthermore depend on the phenotype of focal individuals (Hyp. 1.b, figure 1), we specifically focused here on the phenotype dependence of social information use about food availability. For instance, we expected an increased probability of movements when focal individuals are in low condition and provided with information about a high amount of resources, long-term high habitat quality, or low competition in the vicinity. In contrast to average social information, the contrast of traits between informants might reflect the heterogeneity of information on surrounding environments (Hyp. 1.c, figure 1). Individuals might choose to decrease their movements when heterogeneity of information about surrounding environments, and thus possibly information uncertainty, increases (Riotte-Lambert et al. 2020, e.g. Heinen and Stephens, 2006). Finally, we expected focal individuals to adjust movement direction depending on social information differences between sources (Hyp. 2, figure 1), possibly moving towards the chamber of the informant having access to food, with a better physical condition, displaying low competitive behaviors, or being of the opposite gender.

## Materials & Methods

### Species and study sites

*Zootoca vivipara* (Jacquin 1787) is a small size ground-living species in the Lacertidae family. This widespread species spans Northern Europe and Asia and lives in heathlands, bogs, and wet grasslands. Individuals used in this study have been sampled in seven populations in the Massif Central mountain range (France). These sites range from 1000m to 1500m and cover the diversity of possible habitats in this region (Rutschmann et al. 2016). In the Massif Central, mating takes place just after individuals emerge from hibernation, between March and April. Parturition usually occurs between late June and late July, depending on temperature conditions (Rutschmann et al. 2016). In our sites the current mode of reproduction is ovoviviparity and juveniles emerge from the egg within a few hours after parturition. Some of the juveniles disperse from their natal site a few days after birth (Massot 1992).

### Capture and rearing condition

The capture and rearing conditions have been validated by an ethical committee (DAP number 5897-2018070615164391-v3). Twenty pregnant females were captured at each site between June 12th and 24th, in 2019. These females were brought to a field laboratory, where we measured snout to vent length. Females were maintained in individual plastic terrariums (18.5 × 12 × 11 cm), containing a shelter made from two slots of a cardboard egg-box and a 2 cm substrate of sterilized soil (Massot and Clobert 2000). Terrariums were placed under an incandescent bulb of 25W providing light and heat for 6 hours a day to allow basking (from 9 a.m. to 12 p.m. and from 2 p.m. to 5 p.m.). Terrariums were sprayed with water three times a day. Females were fed with three mealworms, every second day.

After parturition (between July 2nd and 24th), neonates from a same clutch were isolated from their mother in a terrarium (day 0). Females that just gave birth were immediately weighed. Neonates’ snout to vent length (SVL) and body mass (BM) were measured the day after their birth (day 1), before any feeding treatment. Sex was assessed following the Lecomte et al. 1992 method. The same day, neonates were isolated to individual terrariums (25 × 15 × 15 cm), containing a shelter made from two slots of a cardboard egg-box and layered with two sheets of absorbent paper. Juveniles were left for another day (day 2) in their respective terrarium before experiments started on day 3, so they could consider this terrarium as their living area (Aragon et al. 2006 b.). All juveniles and mothers were fed and released at the mother’s capture site on day 4.

### Experimental design

The experiment aimed at testing if the spatial decisions of a focal individual were influenced by informants’ phenotypes and food intake. Each of the 56 replicates of the experiment required three juveniles (two informants and one focal individual). Each juvenile was tested in a single replicate. For each replicate, juveniles were selected among clutches of mothers from the same capture site. When possible, informants had the same laying date. Most experimental replicates (n=37) took place 3 days after the birth of focal individuals but some replicates happened 2- (n=10) or 4-days (n=9) after birth when there were too few births on the same day. Similarly, 2 (n=10) and 4 days old (n=13) informants were used when necessary. Seventeen replicates were associated with a difference of age between the focal individual and at least one of the informants. When possible, the three individuals were selected from different broods, but informants from the same brood were used in the same experimental replicate when there were too few births (n=19).

One day before the confrontation between individuals, one of the two informants had access to food: three small crickets, from 3 to 5 mm, were introduced in the terrarium. The number of consumed crickets (0 to 3) was counted just before the experiment (referred to as the fed informant’s food intake further on). The focal individual was never fed before the experiment. Our experiment further manipulated the combination of informants’ sexes orthogonally to the information about food access. Each focal female (n=29) or male (n=27) was confronted with two informant males, two informant females, or one informant male and one informant female. These combinations were balanced between replicates within an experimental day. In the replicates with one informant male and one informant female, the fed informant was the male in nearly half of the replicates (n=14) and the female in the other replicates (n=16). Furthermore, the three juveniles had variable phenotypes (SVL and BM, mothers’ SVL and BM) that we did not control for in the preparation of our experiments. However, we analyzed their effects on movement decisions, since they displayed sufficient variability to be potential social cues (informants SVL: 20 ± 1 mm, informants BM: 157.7 ± 18.2 mg, informants’ mothers SVL: 62.2 ± 4.1 mm, informants’ mothers BM: 3616 ± 566.7 mg).

Replicates took place from July 5th to 27th between 7 a.m. and 8 p.m. We summed up the timing of experiments in a categorical variable, accounting for the lighting periods in the field laboratory, with four classes: early morning (7 a.m. to 9 a.m., n=13), morning (10 a.m. to 12 a.m, n=23), afternoon (1 p.m. to 4 p.m, n=13) and evening (5 p.m to 8 p.m, n=7).

By-products of experimental constraint (age difference between informants, informant kinship and time windows) had no significant effect on relocation probability when individually added in the selected model about relocation probability (respectively (z-value, p) = (−1.08, 0.28) / (−0.85, 0.39) / (from 0.26 to 1.22, from 0.22 to 0.80)). Difference in age between informants did not impact movement decision when added in the model about movement direction (*χ*^2^ = 1.39, p = 0.24).

### Experimental assay

The home terrarium of the focal individual was placed on an isolated table. Corridors (PVC tubes of 25 cm length and 16 mm internal diameter) were introduced at each side of this terrarium. Informants were placed in corridors’ extremities and their arrival in the focal terrarium was synchronized by slightly brushing their tails (figure S1). We alternated the introduction position (left or right) of the fed informant and males and females between each replicate so that the position was not biased towards a treatment or an informant’s sex. Once the informants entered the focal terrarium, exits were plugged. The dimension of the focal terrarium (25 × 15 × 15 cm) was sufficient to accommodate the three juveniles together while allowing them to avoid each other. The three juveniles interacted together for thirty minutes (figure S1). After that period, informants were put back in their respective terrariums. In the home terrarium, absorbent paper, shelter, and heat/light source were removed to promote departure (Aragon et al., 2006 b.). The focal individual was left five minutes in these conditions to accommodate (figure S1). Then the corridors used previously were attached again at each extremity of the focal terrariums (without any modification since the informants’ passage) and connected to two identical and clean terrariums. The focal individual was left thirty minutes in this system (figure S1). After this time, the experiment was stopped and we observed if the focal individual had left his terrarium and in which direction. Terrariums in which focal individuals arrived were washed with water between replicates. Corridors were used only once. Experiments were entirely filmed with three webcams (Creative Live Camera Sync HD 720p) placed above each terrarium to analyze the three individuals’ joint behaviors.

### Data analyses

#### Joint behaviors of focal and informant individuals

Behaviors of focal and informant juveniles were measured to test their impact on relocation decisions (Phenotype-dependence of movement and Hyp. 1.a, in figure S1). Due to technical constraints (i.e., videos quality), we could not distinguish the behavior of informants and focal individuals. As a consequence, we analyzed the joint behaviors of the three juveniles together through the last twenty minutes of their confrontation, the first ten minutes being considered as an accommodation time (Cote et al. 2007, 2008 a.). We used BORIS software (Friard and Gamba, 2016) to quantify the following behavioral traits (the description and interpretation of behaviors are detailed in table 1): activity, sheltering, escaping attempt, boldness behavior, non-aggressive proximity, competitive interactions, and tongue-flicking. These behaviors were considered apart from other focal individuals’ traits or informants’ traits because they described the behaviors of both focal and informant individuals simultaneously and cannot individually be ascribed.

**Table 1:** Joint behaviors of the focal individuals and informants.

| Behavior | Description | Unit | Mean<br>value | Sd |
| --- | --- | --- | --- | --- |
| Activity | Cumulative time spent moving by the three individuals<br>(Aragon et al. 2006 a., Cote et al. 2008 a.) | Seconds | 949 | 389 |
| Sheltering | Cumulative time spent sheltering by the three individuals<br>(Cote et al. 2008 a.) | Seconds | 1381 | 795 |
| Escaping<br>attempt | Time spent by at least one individual trying to escape the<br>terrarium by scratching or climbing the walls (Aragon et<br>al. 2006 a.) | Seconds | 64.8 | 78.6 |
| Boldness | Time spent by at least by one individual basking above the<br>shelter (Cote et al. 2008 b.) | Seconds | 177 | 232 |
| Non-aggressive<br>proximity | Time spent motionless by at least two individuals in close<br>proximity to each other, i.e. at a distance less than the<br>approximate size of an individual (Aragon et al. 2006 b.) | Seconds | 192 | 160 |
| Competitive<br>interaction | Number of contacts between two individuals leading to<br>the flight of at least one individual (Aragon et al. 2006 b.) | Events<br>count | 3.61 | 3.86 |
| Tongue-Flicking | Olfactory cues acquisition that could be interpreted as a<br>chemical exploration (Cooper 1994, Aragon et al. 2006 b.) | Events<br>count | 2.88 | 3.43 |

Description of the joint behaviors of the three juveniles, measured during the last twenty minutes of their confrontation (see Mat&Met). We did not use cumulative times for escaping attempts and boldness behavior because these behaviors were performed extremely rarely by several individuals at the same time (4% and 2.7% respectively). Bibliographic references used to define these metrics are cited in the description column. Mean and standard deviation have been computed over all experimental replicates.

We synthesized behavioral information using a principal component analysis (PCA) (figure S3; all analyses were performed with the “FactoMineR” R package, Lê et al. 2008). Given the explained variance distribution (figure S2-A, computed from eigenvalues, with the same distribution) we retained the first axis, explaining 50.4% of the variance, for subsequent analyses. This axis could be described as the global activity level, with non-negligible (loadings > 0.4, variables close to this threshold were also displayed) positive contribution of activity, competitive interactions, non-aggressive proximity and tongue-flicking behaviors, and a negative contribution of sheltering behavior (figure S2-B and table 2 for details). This axis will be subsequently referred to as juveniles’ joint activity level. The second PCA axis (which mainly described the boldness behavior, figure S2-B) explained a non-negligible part of the variance (18.5%). Yet, when including this variable in our models we found no significant effects of this variable on focal individuals’ relocation (z-value = -1.13, p = 0.26) and it did not change the other variables’ significance.

**Table 2:**
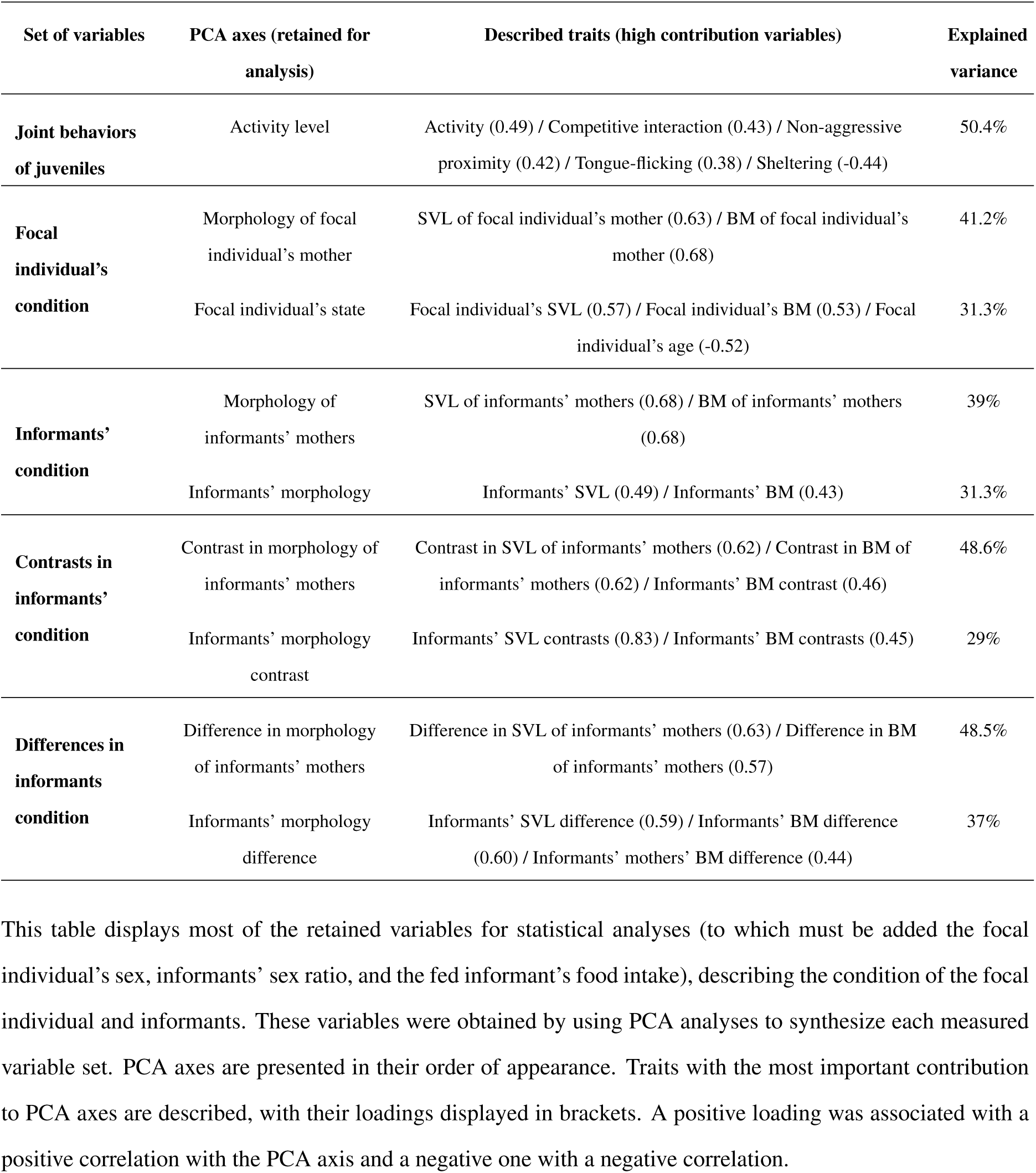
Variables describing the condition of juveniles.

| Set of variables | PCA axes (retained for analysis) | Described traits (high contribution variables) | Explained variance |
| --- | --- | --- | --- |
| **Joint behaviors of juveniles** | Activity level | Activity (0.49) / Competitive interaction (0.43) / Non-aggressive proximity (0.42) / Tongue-flicking (0.38) / Sheltering (-0.44) | 50.4% |
| **Focal individual's condition** | Morphology of focal individual's mother | SVL of focal individual's mother (0.63) / BM of focal individual's mother (0.68) | 41.2% |
|  | Focal individual's state | Focal individual's SVL (0.57) / Focal individual's BM (0.53) / Focal individual's age (-0.52) | 31.3% |
| **Informants' condition** | Morphology of informants' mothers | SVL of informants' mothers (0.68) / BM of informants' mothers (0.68) | 39% |
|  | Informants' morphology | Informants' SVL (0.49) / Informants' BM (0.43) | 31.3% |
| **Contrasts in informants' condition** | Contrast in morphology of informants' mothers | Contrast in SVL of informants' mothers (0.62) / Contrast in BM of informants' mothers (0.62) / Informants' BM contrast (0.46) | 48.6% |
|  | Informants' morphology contrast | Informants' SVL contrasts (0.83) / Informants' BM contrasts (0.45) | 29% |
| **Differences in informants condition** | Difference in morphology of informants' mothers | Difference in SVL of informants' mothers (0.63) / Difference in BM of informants' mothers (0.57) | 48.5% |
|  | Informants' morphology difference | Informants' SVL difference (0.59) / Informants' BM difference (0.60) / Informants' mothers' BM difference (0.44) | 37% |

This table displays most of the retained variables for statistical analyses (to which must be added the focal individual's sex, informants' sex ratio, and the fed informant's food intake), describing the condition of the focal individual and informants. These variables were obtained by using PCA analyses to synthesize each measured variable set. PCA axes are presented in their order of appearance. Traits with the most important contribution to PCA axes are described, with their loadings displayed in brackets. A positive loading was associated with a positive correlation with the PCA axis and a negative one with a negative correlation.

#### Condition of informants and focal individuals

In our models, we considered three groups of variables describing informants’ and focal individuals’ condition (Table 2): the condition of focal individuals (age, SVL, BM, mother’s SVL and BM; testing for phenotype-dependence of movement and Hyp. 1.b in figure 1), the condition of informants (mean age, mean SVL and BM, mean mothers’ SVL and BM; testing for Hyp 1.a and Hyp 1.b in figure 1) and the absolute difference between informants’ traits (SVL, BM, mothers’ SVL and BM; testing for Hyp 1.c in figure 1). Note that for the orientation decision (see later), the latter group of variables was replaced by the raw difference between informants’ traits (SVL, BM, mothers’ SVL and BM, left informant minus right informant traits; testing for Hyp 2 in figure 1), to spatially polarize the contrast between informants. To synthesize these variables we used a PCA and conserved the axes explaining most of the variance in each group (all selected axes explained > 70% of the variance, figures S3-A, S4- A, S5-A, and S6-A). All selected PCA axes and the respective part of explained variance are described in table 2.

Briefly, the first two axes of the focal individual’s condition PCA (figure S3-B) resume the morphology (i.e., SVL and BM concomitant variations) of the focal individual’s mother and the focal individual’s state (define as SVL, BM, and age concomitant variations). The first two axes of the informants’ condition PCA (figure S4-B) resume the morphology of informants’ mothers and informants’ morphology. The first two axes of the contrasts in informants’ condition PCA (figure S5-B) resumed the contrast in the morphology of informants’ mothers and the contrast in informants’ morphology. Finally, the first two axes of the PCA on differences in informants’ condition (figure S6-B) synthesized the difference in morphology of informants’ mothers and difference in informants’ morphology. In addition to these four PCAs axes, we also considered in our models the focal individual’s sex, informants’ sex ratio, and the fed informant’s food intake. Of note, there was no correlation between the fed informant’s food intake and informants’ morphology (Pearson correlation test, p = 0.54) or morphology of informants’ mothers (Pearson correlation test, p = 0.48), and between sex and physical condition (Wilcoxon tests; focal individual’s sex and state: p = 0.47, informants’ sex-ratio and morphology: p= 0.42).

#### Statistical Analyses

All statistical analyses were performed with R software (R Development Core Team, 2008, version 3.6.3). Graphs were produced using the package “ggplot2” (Wickham 2016). First, we analyzed the relocation probability of focal individuals after their confrontation with the two informants (Hyp. 1.a.b.c, figure 1). To do so, we used a Firth logistic regression (Firth 1993, ‘brglm’ function with a logit link, in “brglm” R package, Kosmidis and Firth 2021), a penalized likelihood regression method. This method was chosen to take into account data separation (Heinze and Schemper 2002), i.e. a predictor variable perfectly predicting the outcome variable (Albert and Anderson 1984), and quasi-separated data, that is likely present in our dataset given our sample size. We first tested for the population of origin as a potential random effect (Zuur et al. 2009). Note here that a daily effect was partially nested in the population variable as the different capture sites are associated with different hatching periods (Rutschmann et al., 2016) and as only one or two capture sites were used each day of experiments. The population random effect appeared non-significant (analysis of deviance test between null models with and without random effects, using standard logistic regressions; p = 0.47) and was dropped in our subsequent models. Then, we performed a model selection (Table S1) among the set of candidate variables, describing the informants and focal individual joint behaviors (activity level PCA axis; to test for phenotype-dependence of movement and Hyp. 1.a), the focal individual’s condition (morphology of focal individuals’ mothers and focal individuals’ state PCA axes, and focal individuals’ sex; to test for phenotype-dependence of movement), informants’ condition (morphology of informants’ mothers and informants’ morphology PCA axes, informants’ sex-ratio and the fed informant’s food intake; to answer Hyp. 1.a), contrasts in informants’ condition (contrast in morphology of informants’ mothers and contrast in informants’ morphology PCA axes; to test for Hyp. 1.c) and the interactions between the fed informant’s food intake and the focal individual condition variables (food intake with every three focal individuals’ condition variables; to test for Hyp. 1.b). All used quantitative variables were scaled. This model selection was performed using the function ‘dredge’ (package MuMIn, Barton et al. 2009). Only one model appeared in Δ*AICc <* 2 (threshold for the best models; Burnham and Anderson 2004), this model was used for all subsequent analyses.

The resulting model showed a sufficiently low variance inflation factor (maximal VIF of 1.71) for the interpretation of our statistical results (O’brien 2007). We measured the quality of our selected model by implementing a Nagelkerke pseudo-R-squared (Nagelkerke 1991). Effects of retained variables were tested through Wald tests on the selected model variables (since model comparison approach on Firth’s regression still an on-going research, Kosmidis and Firth 2021). We also used partial Nagelkerke pseudo-R-squared on our model to rank variables by their level of explained variance and relative importance obtained from the model selection (sum of Akaike weights) to estimate all the variables’ degree of importance (including the ones that did not appear in our selected model).

A second analysis was conducted to test which of the informants’ traits influenced the direction of relocation (Hyp. 2, figure 1) when focal individuals left their terrarium (n=22). This time, we used simple logistic regressions with a binary response variable (leave toward left or right). In this model, we used the spatial distribution of male and female informants (female coming from the right and male coming from the left, male coming from the right and female coming from the left or the same sex left and right), fed informant spatial origin (coming from left or right), difference in informants’ morphology (Table 2) and difference in morphology of informants’ mothers (Table 2). The model was diagnosed as presented before, with an analysis of deviance instead of Wald tests for testing the variables effects (likelihood-ratio tests, “car” R package, Fox and Weisberg 2018). Again, we obtained a sufficiently low VIF (maximum equal to 1.38) for the interpretation of our results.

## Results

Over the 56 experimental replicates, 22 focal individuals left their terrariums. The regression results revealed significant (p<0.05), or marginally significant (0.05<p<0.1), effects of focal individuals’ state (age, morphology) and sex, informants’ traits (maternal morphology, the fed informant’s food intake), focal individuals’ and informants’ joint activity level, and the interaction term between the focal individual’s state and the fed informant’s food intake. All tests’ statistics are available in Table 3. Shortly, the relocation probability of focal individuals decreased with their age and increased with their morphology (figure S7-A) and was lower for males compared to females (figure S7-B). The relocation probability further decreased with larger informants’ maternal morphology (figure S8-B, table 3) and the fed informant’s food intake (figure S8-A, table 3). Relocation also depended on the joint activity level of focal and informant individuals (figure S9): it increased with the increase of activity, with higher levels of social interaction (non-aggressive proximity and competitive interactions), and with higher levels of exploration behaviors but decreased with increased sheltering behaviors (Mat&Met, figure S2-B and Table 2). We also found that the food intake of the fed informant interacted with the phenotype of the focal individual to impact the focal individual’s probability of relocation (figure 2): relocation probability increased for an informant with poor food intake confronted to a focal individual with a low score of individual’s state (i.e. smaller morphological traits and older age) or for an informant with high food intake confronted to a focal individual with a high score of individual’s state (i.e. larger morphological traits and younger age). Overall, the fitting quality of this model was good, with a Nagelkerke R-squared of 0.68.

**Table 3:**
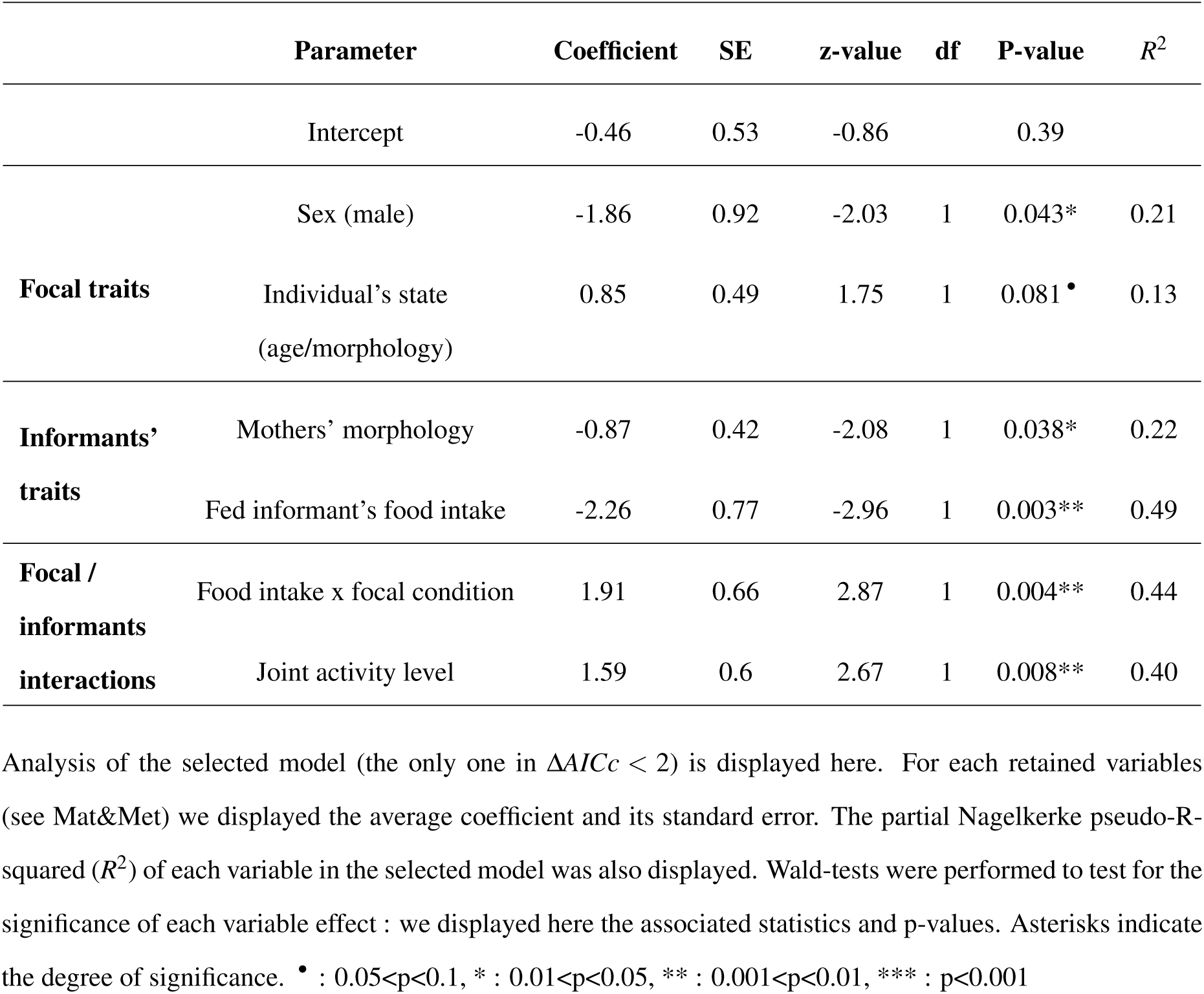
Selected Firth’s logistic regression about focal individuals’ relocation probability.

**Figure 2:**
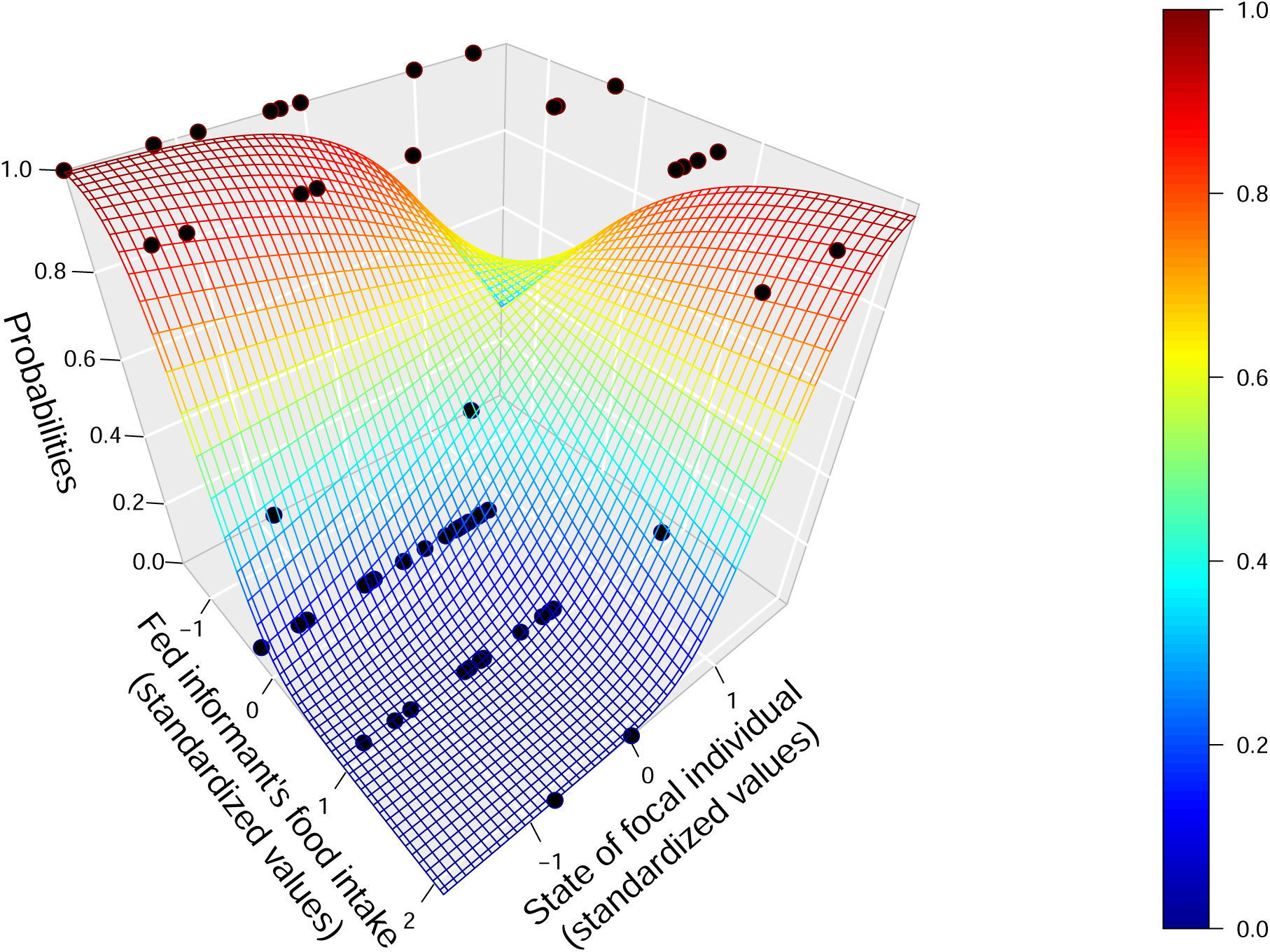
Joint effects of the fed informant’s food intake and the focal individual’s state on relocation probability of focal individuals. We looked at the distribution of focal individuals’ relocation predicted probability as a function of the fed informant’s food intake (number of eaten crickets, standardized values) and the focal individual’s state (standardized values). The graph was produced using the Firth’s logistic regression results (see Table 3), by plotting the predicted probabilities as a function of the variable of interests’ and the intercept’s coefficients (all other coefficients were fixed to 0, i.e. their average or their baseline level as they are standardized). Black dots display observations from all experimental replicates: a dot on the 0% probability surface corresponds to a focal individual who did not leave his terrarium, a dot on the 100% probability surface corresponds to a focal individual who left his terrarium.

We computed explained variance estimates for each variable included in the selected model by using Nagelkerke’s partial R-squared (Table 3). Food intake of fed informants and joint behaviors had relatively high partial r-squared of 0.49 and 0.4; the sex of focal individuals and morphology of informants’ mothers had respective r-squared of 0.21 and 0.22; the state of focal individuals (age and morphology) had an r-squared of 0.13. The r-squared of the interaction between the fed informants’ food intake and the focal individual’s state (0.44) was rather important relatively to the previously described values. The relative importance of all tested variables (obtained from model selection, Table S1) showed that the importance of the other non-selected variables was much lower (relative importance inferior to 0.27 for non-selected variables and superior to 0.84 for selected variables).

We then analyzed the movement direction of the 22 focal individuals which left their terrariums; nine individuals went to the right and thirteen to the left. No significant effect was found among the tested variables, including the feeding treatment (Table 4, figure S10). Overall, we obtained a Nagelkerke R-squared of 0.15, suggesting a poor quality for the model.

**Table 4:** Logistic regression on direction taking (analysis of deviance).

| Parameter | $\chi^2$ | df | P-value |
| --- | --- | --- | --- |
| Fed informant origin | 0.71 | 1 | 0.4 |
| Sex contrast between informants | 1.88 | 2 | 0.39 |
| Morphologies contrast between informants | 0.004 | 1 | 0.95 |
| Morphologies contrast between informants' mothers | 0.034 | 1 | 0.85 |

Results of the logistic regression on focal individuals' direction taking are displayed here. An analysis of deviance was performed to test for the significance of each variable effect (likelihood ratio tests). For each test, the chi-squared statistic and the associated p-values are displayed. Asterisks indicate the degree of significance.
• : $0.05 < p < 0.1$ , \* : $0.01 < p < 0.05$ , \*\* : $0.001 < p < 0.01$ , \*\*\* : $p < 0.001$

## Discussion

We experimentally investigated how social information is used for movement decisions when simultaneous sources of information (i.e., informant individuals), coming from different environments, are available for decision making.

We found the relocation probability of focal individuals to be phenotype-dependent (in support of the expected phenotype-dependence of movement): relocation was more likely to occur for females, to decrease with individual’s age, and to increase with their score of morphology (SVL and BM). We also found that the relocation probability of focal individuals increased when informants ate fewer available prey and originated from mothers with smaller morphological traits (low BM and SVL), suggesting the use of averaged social information for movement decisions (in support of Hyp. 1.a: figure 1). We further observed a phenotype-dependence of the use of social information about food availability (in support of Hyp. 1.b: figure 1). We found no significant influence of contrasts in traits between informants, potentially representative of information variability or reliability, on the relocation probability of focal individuals (contrary to the expectations from Hyp. 1.c: figure 1). Finally, when focal individuals left their terrarium, the position of the informant having access to food did not influence the direction of the movement, as for other differences in traits between informants (contrary to the expectations from Hyp. 2: figure 1).

### Phenotype dependence of relocation

In many species, movement from one location to another can be related to environmental factors or the individual’s phenotype (respectively context- and phenotype-dependence, Bowler and Benton, 2005). In the common lizard for example, dispersal is known to depend on the juvenile’s phenotype such as its level of stress or its physical condition (Clobert et al., 2012). In our experiments, we further observed that the focal individual’s state variable correlated with the relocation probability: it increased with larger morphology (BM and SVL) and decreased with age. The morphology of a neonate lizard, just after birth, directly reflects the amount of energetic reserve available from initial yolk reserves in the egg and therefore influences its performance in the early stage of life (Sinervo 1990, Olsson et al. 2002). As movements implied energetic cost (for displacement itself or potential interactions with competitors and predators, Bonte et al. 2012), larger reserves should provide an advantage for successfully relocating toward another habitat if necessary. Such a relationship between juveniles’ physical condition and movements had already been observed in common lizards for natal dispersal (Meylan et al. 2002). Similarly, in the absence of any food intake, neonates only rely on the energetic reserves inherited from the eggs’ yolk for early physiological performance. As a consequence, younger individuals may be more inclined to allocate such reserves towards relocation than older individuals, forced to conserve their energy for foraging. It could be an advantage to rapidly use their natal reserves for displacement or exploration, especially as early-stage appeared to be central for future survival (Mugabo et al. 2010, Massot and Aragon, 2013). Another possible hypothesis to explain the increase of relocation probability for younger juveniles is that individuals may try to escape competition with their mother and sibling by relocating as soon as possible from their birth location (Galliard et al. 2003, Cote et al. 2007, Cote and Clobert 2010). We also observed an effect of focal individuals’ sex on their relocation probability: the propensity for juveniles to relocate was higher for females than for males. This is a surprising result as male-biased movements are often observed in lizard species (e.g., Doughty et al. 1994, Schofield et al. 2012), including the common lizard (Galliard et al. 2005 b.). Yet, this result echoes what has been found in earlier experiments (Aragon et al. 2006 b.), where, when relocating after a confrontation with another neonate, female juveniles relocated with lower latency than males. Thus, it is likely that the observed female-biased movement would have disappeared if we had let the juveniles dispersed for a longer period, to potentially give way to male-biased movements.

Finally, the positive effect of the joint level of activity of informants and focal individual on the focal individual’s relocation probability might result from independent or joint effects of the focal individual behavior or of informants’ competitiveness (that we will discuss later) that we cannot distinguish in our experiment. When considering the focal individual’s behavior, a more active and explorative individual (high movement rate, low sheltering, high chemical exploration through tongue-flicking, high non-aggressive proximity with conspecifics) is indeed more likely to leave its home environment (Cote et al. 2010).

### Use of social information from multiple sources

In our experiment, interacting conspecifics might convey information about either the local or the distant habitat. Although distinguishing between these two non-exclusive hypotheses would require further experimental investigation that is beyond the scope of this paper, we here elaborate on the patterns expected under each scenario and how they match (or not) with our findings. If informants mainly carried information about the local habitat, we would expect focal individuals to stay in the current environments when confronting individuals bearing cues or signals related to high-quality environments. On the contrary, a relocation of the focal individual in presence of individuals carrying cues or signals related to high-quality environments would suggest that the phenotypic traits of informants convey information about their habitat of origin.

We observed that relocation propensity increased when the morphology of informants’ mothers decreased, and when focal lizards were facing fed informants which did not eat much. These observations are in favor of our second hypothesis as they are indicators of low resource availability in the close environment, and therefore related to an avoidance of a poor local environment given the importance of resources for survival (Mugabo et al. 2010, 2011, Massot and Aragon 2013). The physical condition of informants’ mothers could also be representative more broadly of habitat quality as it could vary with other important environmental parameters as density (Massot et al. 1992) or abiotic parameters as temperature (Chamaillé-Jammes et al. 2006). Avoidance of informants whose mothers were of poor physical condition could then reflect the avoidance of a local environment of low quality. Such an avoidance had already been observed with dispersal increase in case of too high competition (Léna et al. 1998, Cote et al. 2008 a.) or abiotic parameter that does not match energetic requirement (Bestion et al. 2015). We also observed that relocation propensity increased with juveniles’ (informants and focal individual) activity. More knowledge about the individual behaviors of informants would have helped to refine our interpretations but the presence of active and aggressive conspecifics is likely to reflect high levels of direct competition in the local environment (Sih et al. 2004, e.g. Garland et al. 1990). A measure of stress hormones (as corticosterone, see Belliure et al. 2004) would be useful to precise the influence of informants’ stress level on focal individuals’ relocation.

Finally, we also observed that relocation probability increased when the fed informant food intake and the focal individual’s physical condition were both low or both high. Again, this result suggests that relocation preferentially occurs when local resource availability does not match the focal individual phenotype. In the first case, individuals seemed to avoid a local environment with insufficient resources given their conditions (low energetic reserves), while in the second case, they seemed to escape unnecessary competition for resources while having a good enough physical condition to relocate toward a less competitive habitat. Such results further suggest that the use of social information is mediated by the phenotype of the focal individuals, a dependency that has been already observed in previous studies (Cote and Clobert 2007, Cote and Clobert 2010, but also Parejo et al. 2007 in other species) but never with information on direct resources availability at stake.

### On the meaning of informants’ traits

In our experiment, interacting conspecifics might convey information about either the local or the distant habitat. Although distinguishing between these two non-exclusive hypotheses would require further experimental investigation that is beyond the scope of this paper, we here elaborate on the patterns expected under each scenario and how they match (or not) with our findings. If informants mainly carried information about the local habitat, we would expect focal individuals to stay in the current environments when confronting individuals bearing cues or signals related to high-quality environments. On the contrary, a relocation of the focal individual in presence of individuals carrying cues or signals related to high-quality environments would suggest that the phenotypic traits of informants convey information about their habitat of origin.

We observed that relocation propensity increased when the morphology of informants’ mothers decreased, and when focal lizards were facing fed informants which did not eat much. These observations are in favor of our second hypothesis as they are indicators of low resource availability in the close environment, and therefore related to an avoidance of a poor local environment given the importance of resources for survival (Mugabo et al. 2010, 2011, Massot and Aragon 2013). The physical condition of informants’ mothers could also be representative more broadly of habitat quality as it could vary with other important environmental parameters as density (Massot et al. 1992) or abiotic parameters as temperature (Chamaillé-Jammes et al. 2006). Avoidance of informants whose mothers were of poor physical condition could then reflect the avoidance of a local environment of low quality. Such an avoidance had already been observed with dispersal increase in case of too high competition (Léna et al. 1998, Cote et al. 2008 a.) or abiotic parameter that does not match energetic requirement (Bestion et al. 2015). Yet, we also observed that relocation propensity increased with juveniles’ (informants and focal individual) activity. More knowledge about the individual behaviors of informants would have helped to refine our interpretations but the presence of active and aggressive conspecifics is likely to reflect high levels of direct competition in the local environment (Sih et al. 2004, e.g. Garland et al. 1990). A measure of stress hormones (as corticosterone, see Belliure et al. 2004) would be useful to precise the influence of informants’ stress level on focal individuals’ relocation.

Finally, we also observed that relocation probability increased when the fed informant food intake and the focal individual’s physical condition were both low or both high. Again, this result suggests that relocation preferentially occurs when local resource availability does not match the focal individual phenotype. In the first case, individuals seemed to avoid a local environment with insufficient resources given their conditions (low energetic reserves), while in the second case, they seemed to escape unnecessary competition for resources while having a good enough physical condition to relocate toward a less competitive habitat. Such results further suggest that the use of social information is mediated by the phenotype of the focal individuals, a phenotype dependence of social information use that has been observed previously (Cote and Clobert 2007, Cote and Clobert 2010, but also Parejo et al. 2007 in other species) but never with information on direct resources availability at stake.

### Direction of movement

Previous findings have shown a limited but existing ability to orientate in space for the common lizard (Strijbosch et al, 1983). Here, we found no effect of the difference in food access between informants or other observed contrasts between informants on movement orientation when relocation occurred. Given the small sample size for direction analyses (22 replicates), we have to be very cautious about the validity of such effects. These results might suggest that focal individuals considered social cues or signals from present information sources as information about local conditions, for which no orientation is needed. Alternatively, individuals might not have had access to sufficient cues for visual orientation, the design being symmetrical and the arrival lasting few seconds only. Decisions on direction would then mainly rely on informants’ arrival with olfactory signals or cues left by informants in corridors (Aragon et al. 2006 a. and c.). Further experiments, focusing for example on pheromones carried by informants, would be necessary to make any conclusion on the actual use of these odors.

## Conclusion

Our experiment showed that the averaged social information coming from multiple sources was transmitted by interacting conspecifics, with mediation in the use of some information by the phenotype of focal individuals. Yet, contrasts between information sources were not used in our experiments for relocation decisions or orientation. The importance of these information transfers for spatial use could be understood as surrounding habitat quality assessment, with indication on resource availability and degree of intra-specific competition. We also showed that, for the common lizard, information prioritization could occur, with a preference over information related to the immediate environment.

## Author contribution

MB, MR, JC, SJ and AR designed the experiment. MB, JC and AR performed the field work. MB performed all experiments and analyses. MB, MR, JC, SJ and AR wrote the manuscript.

## Acknowledgements

We thank S. Liegeois, C. Fosse, A. Le Pajolec and C. Lauden for their help during experiments and field work. We thank the Parc National des Cévennes for allowing us to use the different sampled sites. This work beneficiated from the scientific environment of the Laboratoire d’Excellence entitled TULIP (ANR-10-LABX-41). The ‘Office Nationale des Forêts’, the ‘Parc National des Cévennes’, and the regions Auvergne, Rhône Alpes and Languedoc Roussillon delivered permits to capture and handle lizards (permits 81-17 2013-05; 2013274-0002, 2013/DREAL/259). No conflict of interest has to be declared.

## Data availability

Analyses reported in this article can be reproduced using the data provided by Brevet et al. (2021).

