## Supplementary figures and tables for "Social information use for spatial decision in *Zootoca vivipara*"

### Supplementary Materials

#### Figure legends

**Figure S1: Experimental design.** Experimental design is summed up in this scheme. The main four steps are successively presented. Corridors are represented as filled grey rectangles, terrariums as empty rectangles. Accesses closure is represented by the crossed out signs.

**Figure S2: Juveniles' behaviors PCA graph of variables.**

**A- Explained variance of the PCA components.**

**B- PCA graph of variables (first and second axes).** All used variables are described in Mat&Met, scratching refers to escaping attempts, proximity to non-aggressive proximity, aggression to competitive interactions. Each arrow is associated with a behavior displayed at their extremities. Arrows indicate strength and sense of correlation among variables and between variables and the PCA axes. The axes explained variances are displayed on the x-axis and y-axis (percentages).

**Figure S3: Focal individuals' condition PCA graph of variables.**

**A- Explained variance of the PCA components.**

**B- PCA graph of variables (first and second axes).** Each arrow is associated with a focal individual trait (body mass, snout-to-vent length, age) or a trait of the focal individual's mother (body mass, snout-to-vent length) displayed at their extremities. Arrows indicate strength and sense of correlation among variables and between variables and the PCA axes. The axes explained variances are displayed on the x-axis and y-axis (percentages).

**Figure S4: Informants' condition PCA graph of variables.**

**A- Explained variance of the PCA components.**

**B- PCA graph of variables (first and second axes).** Each arrow is associated with an average informants' trait (body mass, snout-to-vent length, age) or an average trait of informants' mothers (body mass, snout-to-vent length) displayed at their extremities. Arrows indicate strength and sense of correlation among variables and between variables and the PCA axes. The axes explained variances are displayed on the x-axis and y-axis (percentages).

**Figure S5: Contrasts in informants' condition PCA graph of variables.**

**A- Explained variance of the PCA components.**

**B- PCA graph of variables (first and second axes).** Each arrow is associated to a trait contrast (absolute difference) between informants (body mass, snout-to-vent length) or informants' mothers (body mass, snout-to-vent length) displayed at their extremities. Arrows indicate strength and sense of correlation among variables and between variables and the PCA axes. The axes explained variances are displayed on the x-axis and y-axis (percentages).

**Figure S6: Differences in informants' condition PCA graph of variables.**

**A- Explained variance of the PCA components.**

**B- PCA graph of variables (first and second axes).** Each arrow is associated with a trait difference between informants (left coming informant minus right coming informant) or informants' mothers (body mass, snout-to-vent length) displayed at their extremities. Arrows indicate strength and sense of correlation among variables and between variables and the PCA axes. The axes explained variances are displayed on the x-axis and y-axis (percentages).

**Figure S7: Focal individuals' traits effects on their relocation probability.** We looked at the distribution of focal individuals' relocation predicted probability as a function of focal individuals' significant traits. Plots were obtained from the Firth's logistic regression results (Table 3) by plotting the predicted probabilities as a function of the variable of interest's and the intercept's coefficients (all other coefficients were fixed to 0, i.e. their average or their baseline level as they are standardized). Effects degree of significance is displayed in table 3. Black dots display observations from all experimental replicates: a dot around the 0% probability line corresponds to a focal individual who did not leave his terrarium, a dot around the 100% probability line corresponds to a focal individual who left his terrarium. These dots were jittered (vertically for quantitative variables, horizontally for qualitative variables) to gain in readability.

**A- Predicted probabilities of focal individuals' relocation as a function of focal individuals' state.** The variable is defined in Mat&Met (Table 2).

**B- Predicted probabilities of focal individuals' relocation as a function of focal individuals' sex.** Red dots display the effects at each possible level (female or male). On the x-axis, "f" refers to female individuals and "m" refers to male individuals.

**Figure S8: Informants' traits effects on focal individuals' relocation probability.** We looked at the distribution of focal individuals' relocation predicted probability as a function of informants' significant traits. Plots were obtained from the Firth's logistic regression results (Table 3) by plotting the predicted probabilities as a function of the variable of interest's and the intercept's coefficients (all other coefficients were fixed to 0, i.e. their average or their baseline level as they are standardized). Effects degree of significance is displayed in table 3. Black dots display observations from all experimental replicates: a dot around the 0% probability line corresponds to a focal individual who did not leave his terrarium, a dot around the 100% probability line corresponds to a focal individual who left his terrarium. These dots were jittered vertically to gain in readability.

**A- Predicted probabilities of focal individuals' relocation as a function of the fed informant's food intake.** The black dots were slightly jittered horizontally to improve readability.

**B- Predicted probabilities of focal individuals' relocation as a function of the morphology of informants' mothers.** The variable is defined in Mat&Met (Table 2).

**Figure S9: Predicted probabilities of focal individuals' relocation as a function of juveniles' joint activity level.**

We looked at the distribution of focal individuals' relocation predicted probability as a function of juveniles' joint activity level. The variable is defined in Mat&Met (Tables 1 and 2). The plot was obtained from the Firth's logistic regression results (Table 3) by plotting the predicted probabilities as a function of the variable of interest's and the intercept's coefficients (all other coefficients were fixed to 0, i.e. their average or their baseline level as they are standardized). Effects degree of significance is displayed in table 3. Black dots display observations from all experimental replicates: a dot around the 0% probability line corresponds to a focal individual who did not leave his terrarium, a dot around the 100% probability line corresponds to a focal individual who left his terrarium. These dots were jittered vertically to gain in readability.

**Figure S10: Feeding treatment absence of effect on orientation.** We looked at the distribution of focal individuals' relocation direction predicted probability as a function of fed informant original direction (through a logistic regression). Effect degree of significance is displayed in table 4. The graph was obtained by plotting the predicted probabilities as a function of the variable of interest's and the intercept's coefficients (all other coefficients were fixed to 0, i.e. their average, or were fixed to their mean level for categorical variables), the package "ggeffects" was used to produce the plot. Black bars represent 95% confidence intervals for predicted probabilities. Red dots represent observations from all experimental replicates: a dot on the 0% line corresponds to a focal individual who did not leave his terrarium, a dot on the 100% line corresponds to a focal individual who left his terrarium. These dots were horizontally jittered to gain in readability.

### Tables

**Table S1:** Model selection on Firth's logistic regressions (focal individual's relocation).

| **Parameters** | | | | | | | | | | Importance |
|---|---|---|---|---|---|---|---|---|---|---|
| **Focal sex** | + | + | + | + | + | + | + | | + | 0.84 |
| **Focal state** | + | + | + | + | + | + | + | + | + | 0.99 |
| Focal mother's morphology | | | | | | | + | | | 0.22 |
| Informants' sexes | | + | | | | | | | + | 0.22 |
| Informants' morphology | | | | + | | | | | | 0.20 |
| **Morphology of informants' mothers** | + | + | + | + | + | + | + | + | + | 0.88 |
| **Fed informant's food intake** | + | + | + | + | + | + | + | + | + | 1 |
| **Joint activity level** | + | + | + | + | + | + | + | + | + | 0.99 |
| Fed informant's food intake x Focal sex | | | | | | + | | | | 0.15 |
| **Fed informant's food intake x Focal state** | + | + | + | + | + | + | + | + | + | 0.99 |
| Fed informant's food intake x Focal mother's morphology | | | | | | | | | | 0.04 |
| Informants' morphology contrast | | | + | | | | | | + | 0.27 |
| Contrast in morphology of informants' mothers | | | | | + | | | | | 0.19 |
| **Model rank** | 1 | 2 | 3 | 4 | 5 | 6 | 7 | 8 | 9 | |
| AICc | 52 | 54.3 | 54.3 | 54.7 | 54.8 | 54.9 | 54.9 | 55.8 | 56.2 | |
| $\Delta AICc$ | 0 | 2.27 | 2.31 | 2.75 | 2.83 | 2.94 | 2.95 | 3.85 | 4.21 | |
| Akaike weight | 0.181 | 0.058 | 0.057 | 0.046 | 0.044 | 0.042 | 0.041 | 0.026 | 0.022 | |

Results of the comprehensive model selection (performed with the 'dredge' function, "MuMIn" R package) on Firth's logistic regressions on focal individuals' relocation probability are displayed here (with all models in $\Delta AICc < 4$). All variables of interest (see Mat&Met) were included in the global model. These variables (in their order of appearance) could be grouped as focal individuals' traits, informants' traits, juveniles joint behaviors, interaction terms between informants and focal individuals (including interactions between the focal individual's traits and food intake of the fed informant) and informants' traits contrasts. Models are ranked according to the corrected Akaike information criterion (AICc), AICc differences between the best model and other ones are also displayed. For each model, the selected variables are indicated by a "+" symbol. Variables present in the selected model (only model in $\Delta AICc < 2$) are in bold. The importance (sum of models' Akaike weight) of each variable is given on the left.

### Figures

**Figure S1:** Experimental design.

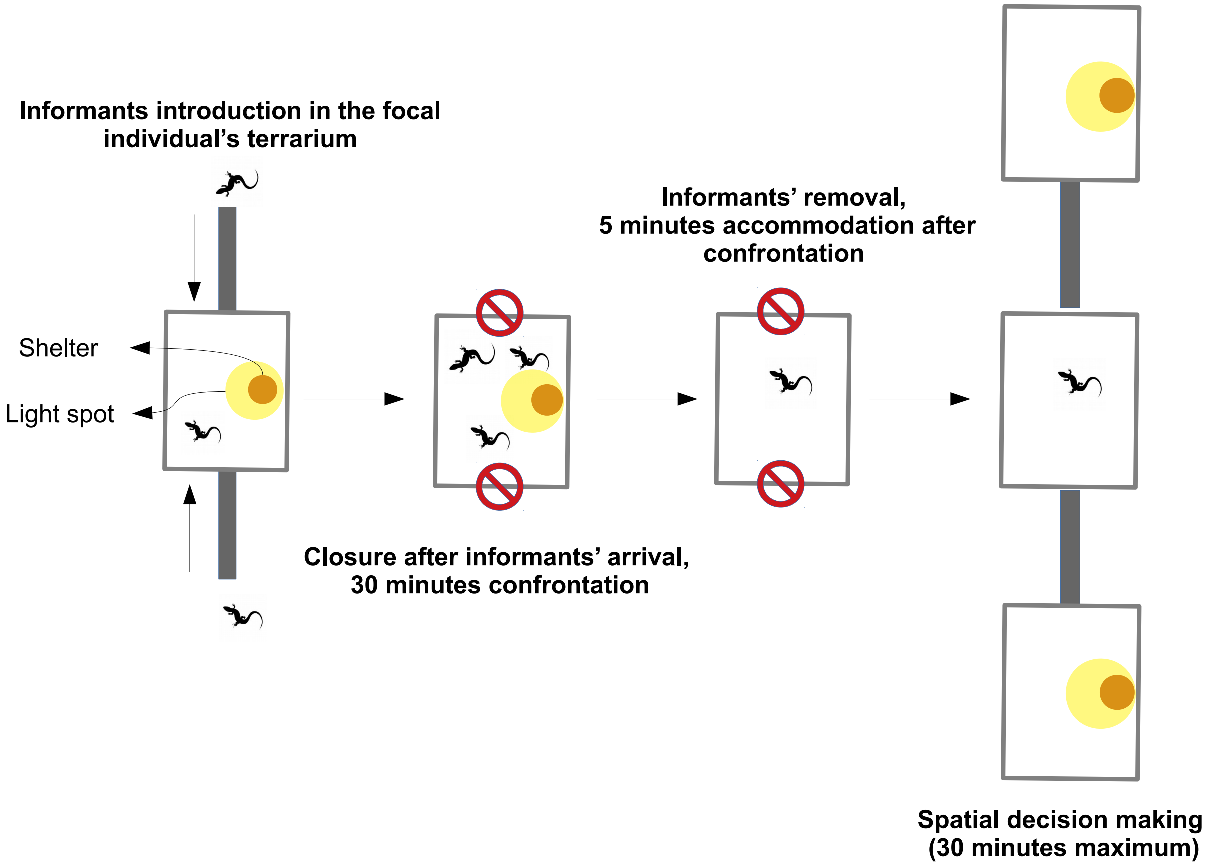

**Figure S2:** Juveniles' behaviors PCA graph of variables.
**A-** Explained variance of the PCA components.
**B-** PCA graph of variables (first and second axes).

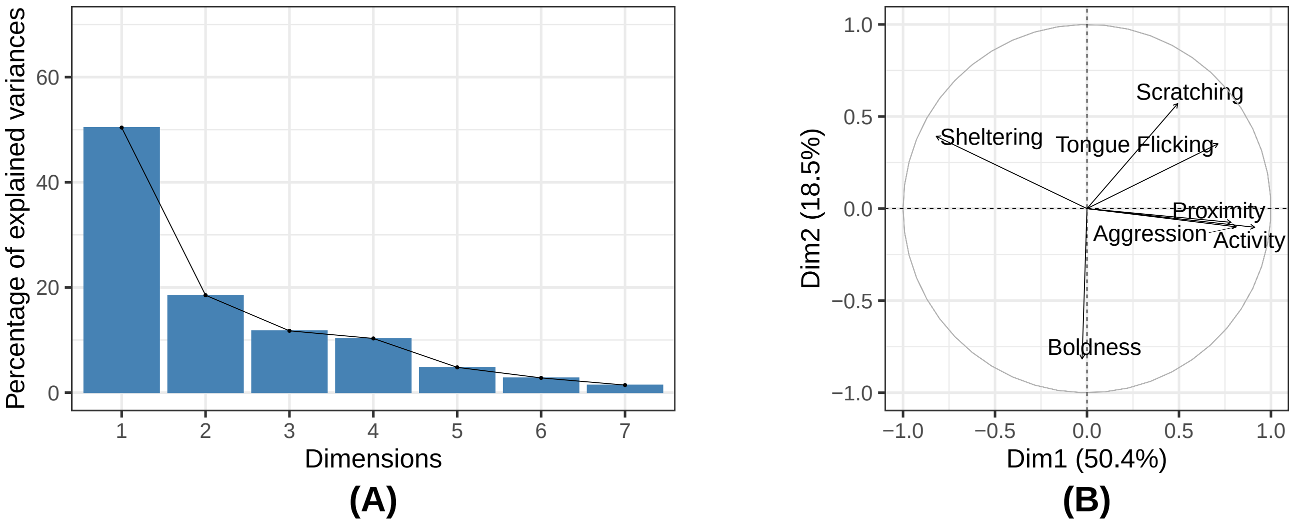


**Figure S3:** Focal individuals' condition PCA graph of variables.
**A-** Explained variance of the PCA components.
**B-** PCA graph of variables (first and second axes).

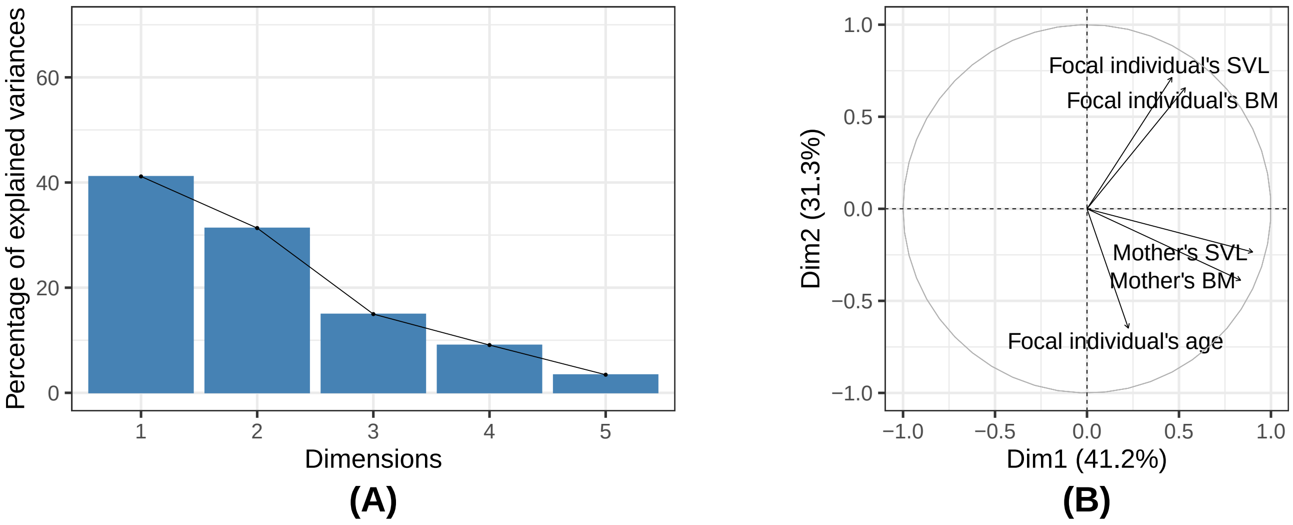

**Figure S4:** Informants' condition PCA graph of variables.
**A-** Explained variance of the PCA components.
**B-** PCA graph of variables (first and second axes).

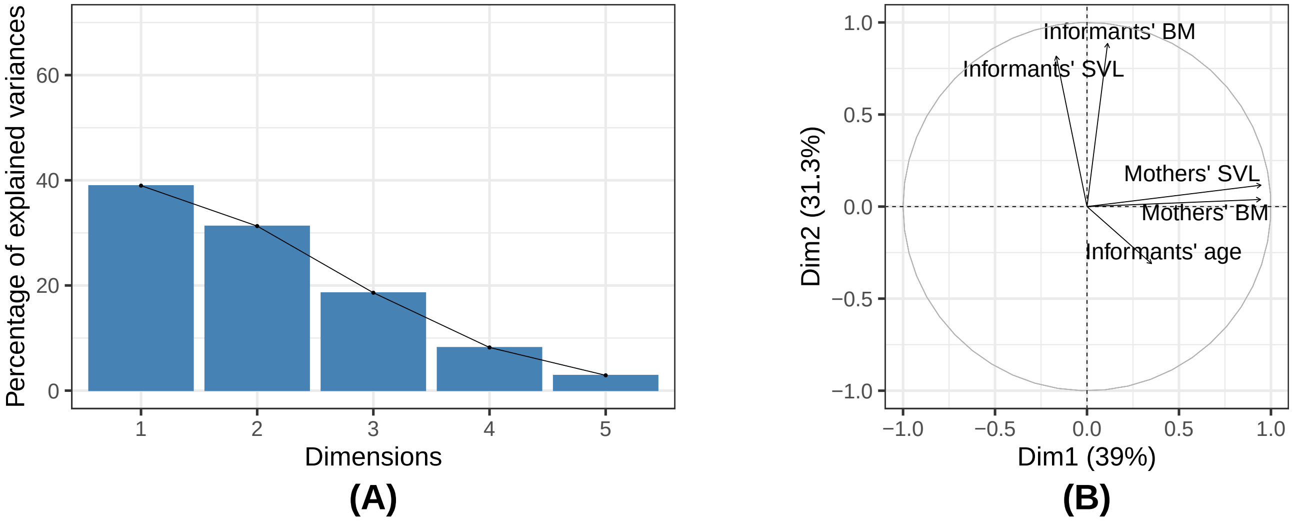


**Figure S5:** Contrasts in informants' condition PCA graph of variables.
**A-** Explained variance of the PCA components.
**B-** PCA graph of variables (first and second axes).

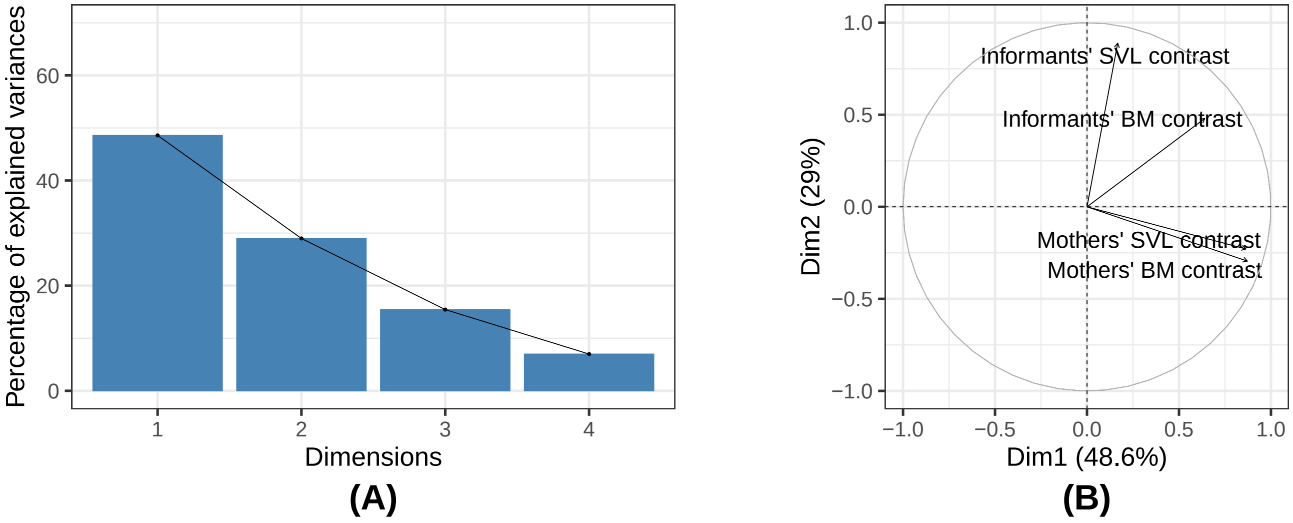

**Figure S6:** Differences in informants' condition PCA graph of variables.
**A-** Explained variance of the PCA components.
**B-** PCA graph of variables (first and second axes).

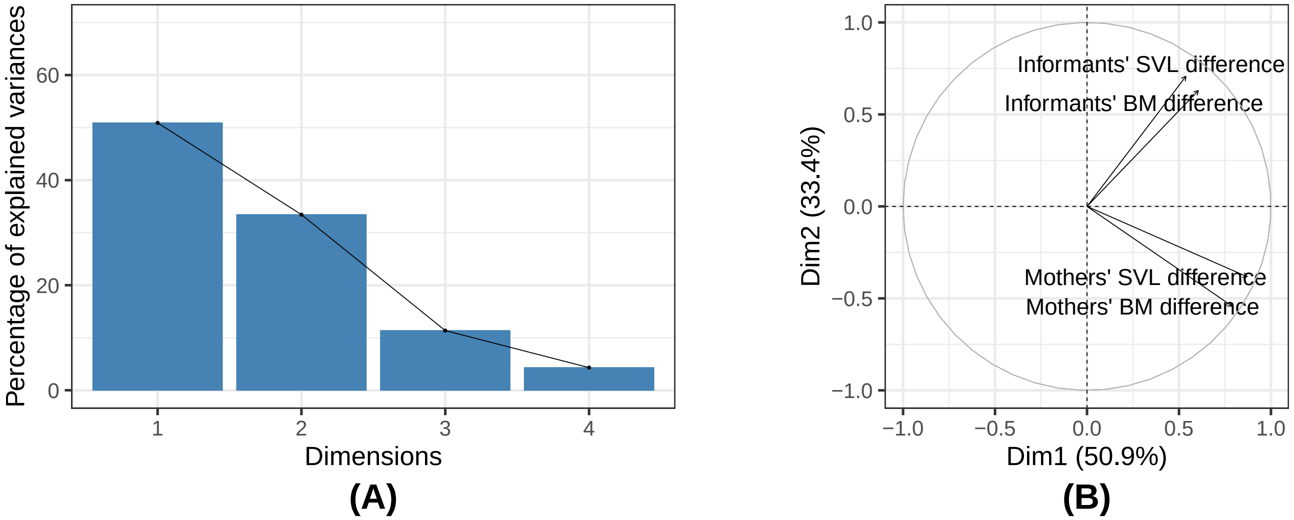


**Figure S7:** Focal individuals' traits effects on their relocation probability.
**A-** Predicted probabilities of focal individuals' relocation as a function of focal individuals' state.
**B-** Predicted probabilities of focal individuals' relocation as a function of focal individuals' sex.

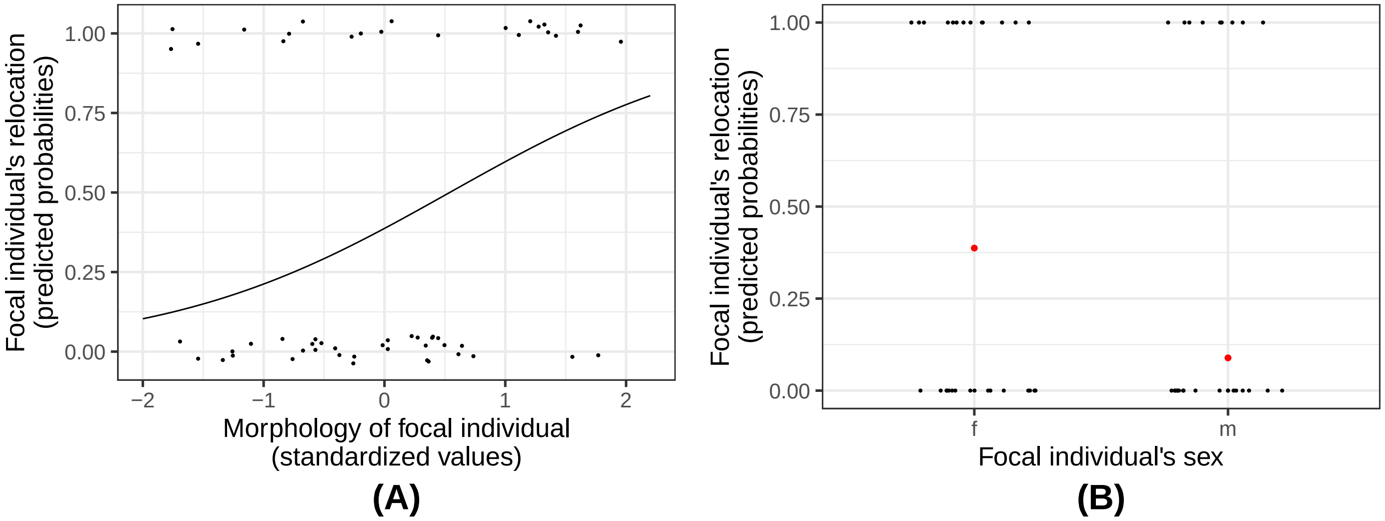

**Figure S8:** Informants' traits effects on focal individuals' relocation probability.
**A-** Predicted probabilities of focal individuals' relocation as a function of the fed informant's food intake.
**B-** Predicted probabilities of focal individuals' relocation as a function of the morphology of informants' mothers.

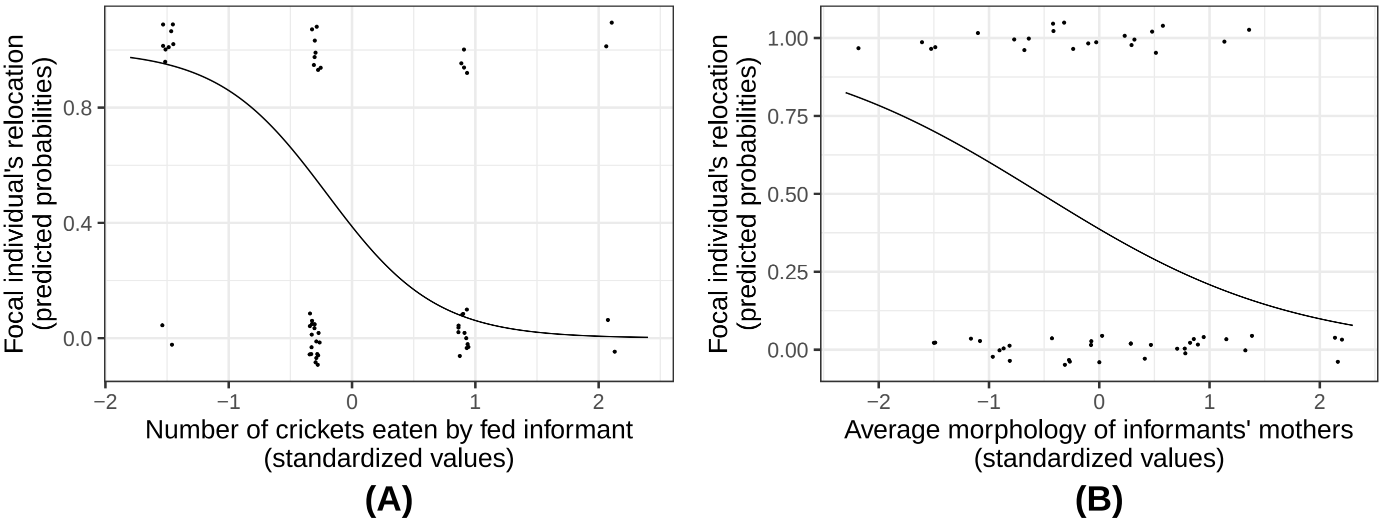

**Figure S9:** Predicted probabilities of focal individuals' relocation as a function of juveniles joint activity level.

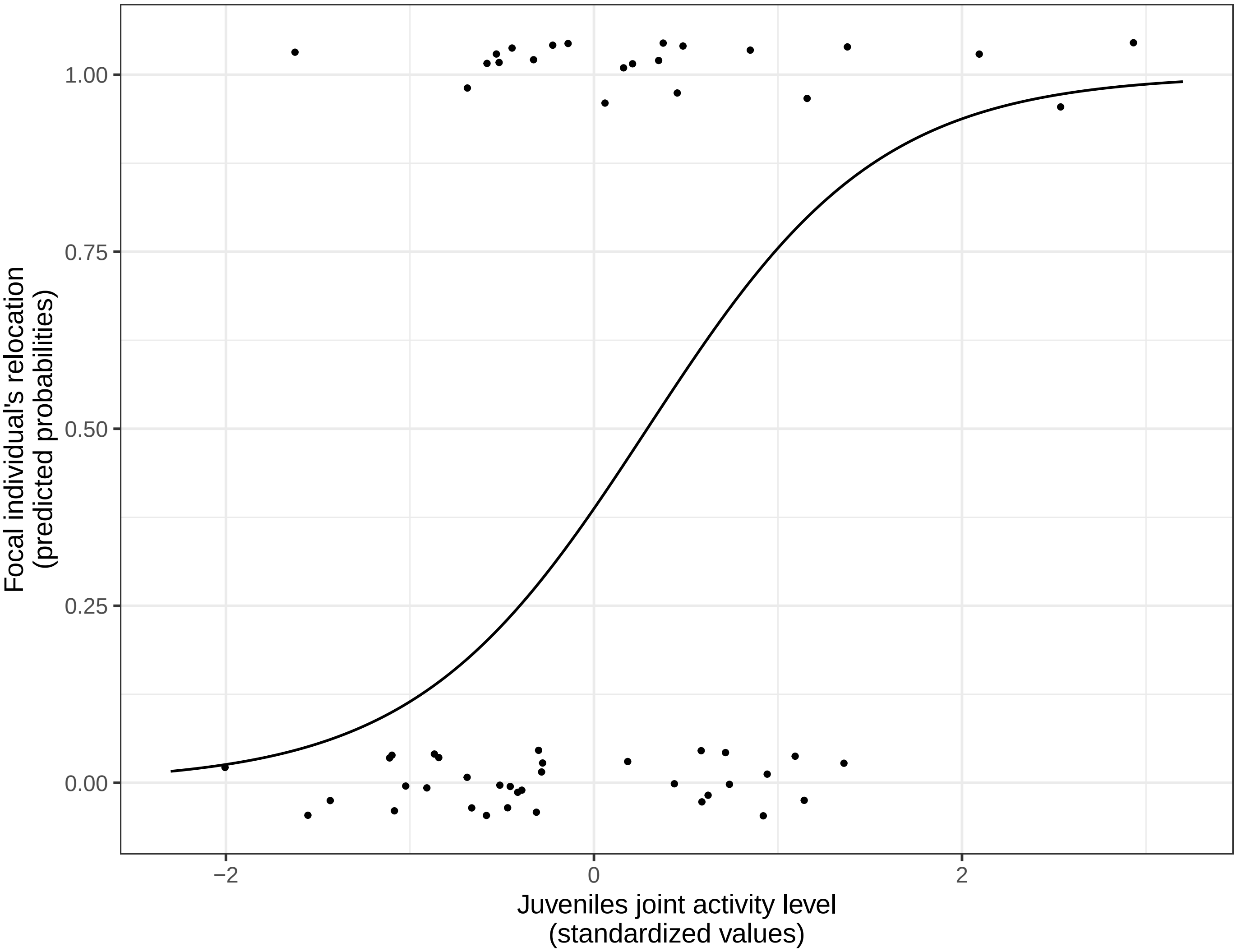

**Figure S10:** Feeding treatment absence of effect on orientation.

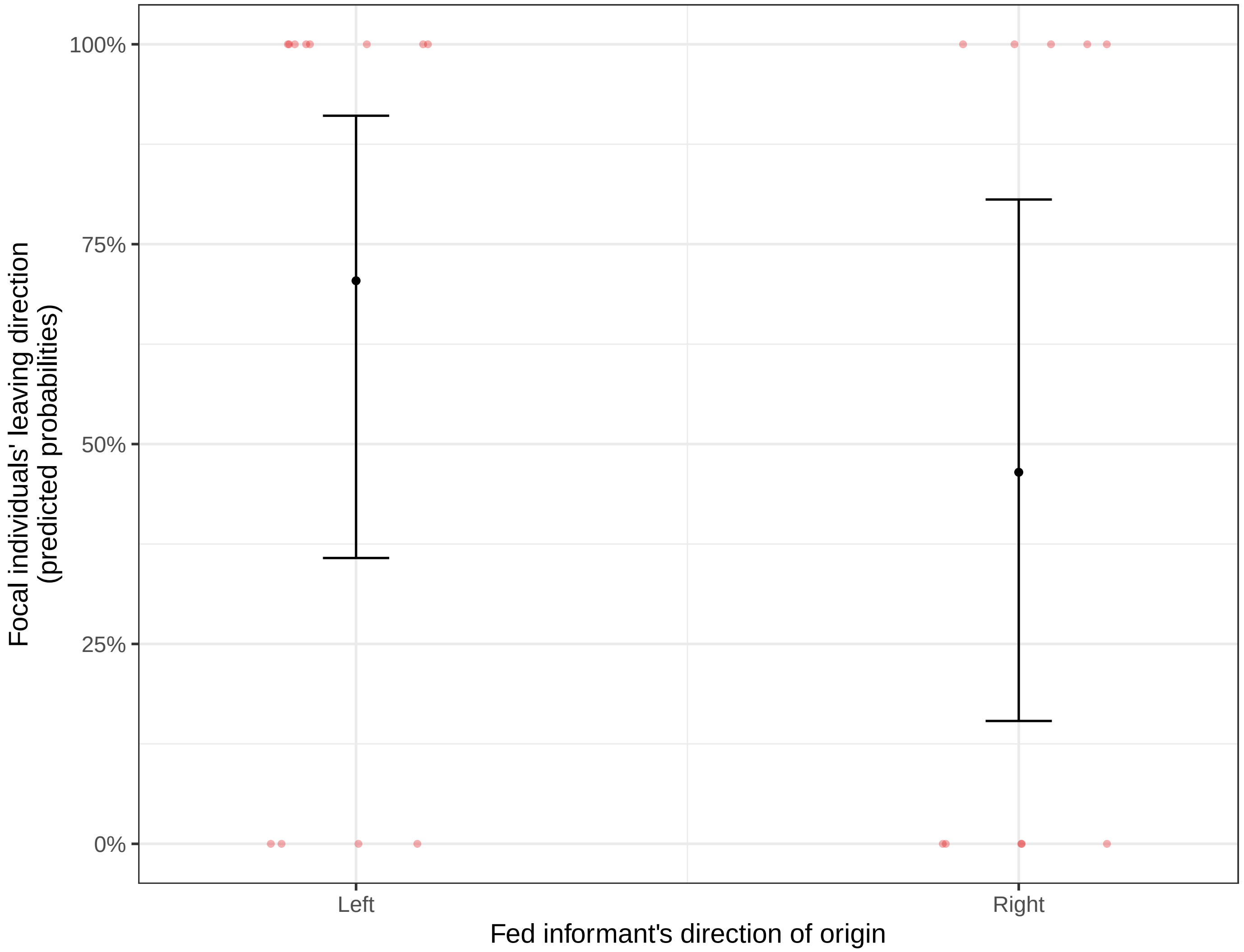
